# Synaptic and Extrasynaptic NMDA Receptors Oppositely Regulate Dendritic Syntaphilin Intrusion in Multiple Sclerosis

**DOI:** 10.64898/2026.07.08.737141

**Authors:** Deepali Mathur, Charlie Zhang, Shing Y Chiu

**Author notes:** Correspondence: Deepali Mathur, PhD Former Research Associate, C/O Professor S.Y. Chiu’s lab, Department of Neuroscience, University of Wisconsin, Wisconsin Institutes for Medical Education and Research (WIMR), 1111 Highland Avenue, Madison, Wisconsin, USA.

## Abstract

Neurodegeneration is a major determinant of disability progression in multiple sclerosis (MS), yet the pathophysiological mechanisms associating inflammation to neuronal insult remain poorly understood. We recently identified Dendritic Syntaphilin Intrusion (DSI), a novel excitoxicity pathway in which the axonal mitochondrial anchor syntaphilin (SNPH) aberrantly translocates into dendrites, causing neurodegeneration in a non-inflammatory model of MS. However, whether this protein intrudes abruptly into dendrites in inflammatory MS pathology is still not clear. Here, we investigated the role of synaptic and extrasynaptic NMDA receptors (NMDAR) in regulating the intrusion of Syntaphilin into dendrites.

Using primary hippocampal neuronal cultures, we examined how the balance between synaptic GluN2A-containing and extrasynaptic GluN2B-containing NMDARs influences DSI under inflammatory conditions. Pharmacological and viral-mediated approaches were employed to manipulate NMDAR subtype activity and evaluate their impact on DSI. Inflammatory cytokines discernibly sensitized neurons to DSI. Our results revealed that blockade of synaptic NMDARs significantly increased DSI, whereas inhibition of extrasynaptic NMDARs reduced DSI. These findings demonstrate opposing roles of NMDAR subtypes, with GluN2A-containing synaptic receptors inhibiting DSI and fostering neuronal survival, while GluN2B-containing extrasynaptic receptors enhancing DSI and neurodegenerative signaling. Manipulation of the GluN2A/GluN2B balance showed opposite effect on DSI, suggesting a relationship between NMDAR subtype signaling and SNPH mislocalization.

Overall, our findings extend the relevance of DSI from non-inflammatory MS to inflammatory MS and identify DSI as a downstream convergence point linking inflammatory cytokines and excitotoxic NMDAR signaling to neuronal insult. These results reveal DSI as a potential mechanistic link between inflammatory signaling and excitotoxic neuronal injury and indicate that modulation of GluN2B-dependent pathways warrants further investigation in inflammatory neurodegenerative disorders.

## Introduction

Multiple sclerosis (MS) is a chronic, inflammatory and an autoimmune disease of the central nervous system characterized by demyelination, and neurodegeneration. Although white matter pathology has traditionally been considered the primary reason of disease progression, increasing evidence indicates that grey matter pathology also contribute to cognitive impairment and irreversible progressive disability independently which is triggered by glutamate excitotoxicity and cytokine exposure (Calabrese et al., 2015; Klaver et al., 2013; Zhang et al., 2021). Importantly, MS pathology cannot be entirely explained by white matter injury alone, suggesting the existence of distinct neurodegenerative mechanisms operating within neuronal compartments (Zhang et al., 2021; Dutta and Trapp, 2011).

Glutamate-mediated excitotoxicity plays a major role in causing neuronal dysfunction and injury (Centonze et al., 2010; Stojanovic et al., 2014; Fairless et al., 2021). Elevated glutamate levels in synaptic region and overactivation of N-methyl-D-aspartate receptors (NMDARs) have been reported in both experimental models and patients with MS, where they contribute to synaptic dysfunction, mitochondrial impairment, and neuronal death (Rossi et al., 2013; Rossi et al., 2014; Fairless et al., 2021). Experimental models of progressive MS have demonstrated substantial grey matter pathology independent of inflammatory demyelination, supporting the existence of distinct neurodegenerative mechanisms (Levy et al., 2010). NMDAR signalling exhibits functional compartmentalization, with synaptic GluN2A-containing receptors generally fostering neuronal survival and plasticity, whereas extra synaptic GluN2B-containing receptors trigger pro-death signalling pathways associated with neurodegeneration (Hardingham and Bading, 2010; Parsons and Raymond, 2014; Zhou et al., 2013; Ge and Wang, 2023). Any disturbance in the balance between synaptic and extra synaptic NMDAR activity has therefore been implicated in several neurodegenerative disorders, including MS (Rossi et al., 2013; Fairless et al., 2021).

Proinflammatory cytokines further augment excitotoxic signalling in MS. In particular, interleukin-1β (IL-1β) facilitates NMDAR-mediated calcium influx by driving Src-family kinases and modifies glutamatergic transmission within cerebellar and cortical circuits (Viviani et al., 2003; Mandolesi et al., 2013; Mandolesi et al., 2015). Cerebellar injury and synaptic pathology are increasingly recognized as important contributors to disability progression in MS, reflecting extensive grey matter involvement beyond classical white matter lesions (Fazio et al., 2008; Wilkins, 2017; Parmar et al., 2018). Damage to cerebellar Purkinje cells caused by glutamate overstimulation is a well-documented phenomenon. It serves as an excellent model to explain how dysregulated glutamatergic pathways contribute to neural cell death in MS (O’Hearn and Molliver, 1993; O’Hearn and Molliver, 1997; Piochon et al., 2007). Furthermore, elevated IL-1β levels correlate with disease progression and promote excitotoxic neurodegeneration through activation of downstream apoptotic pathways (Rossi et al., 2014). These observations suggest that inflammatory factors and NMDAR signaling pathways synergistically promote neuronal degeneration during the progression of MS.

Mitochondrial dysfunction is another hallmark of neurodegeneration in MS and is increasingly recognized as a critical determinant of neuronal loss (Mahad et al., 2015; Devine and Kittler, 2018). Proper mitochondrial distribution in a neuron is regulated by syntaphilin (SNPH), an axonal mitochondrial docking protein that immobilizes mitochondria at sites where energy demand is required (Kang et al., 2008; Chen and Sheng, 2013). We recently identified a novel pathological phenomenon termed Dendritic Syntaphilin Intrusion (DSI), in which SNPH aberrantly translocate from axons into dendrites, leading to mitochondrial mislocalization, dendritic dysfunction, and neurodegeneration (Joshi et al., 2019). Furthermore, deletion of SNPH-mediated mitochondrial anchoring using SNPH-KO protects against neurodegeneration in dysmyelinating models, underscoring the pathological significance of mitochondrial anchoring mechanisms in progressive MS (Joshi et al., 2015).

More recently, we demonstrated that inflammatory cytokines and NMDAR activation interact to trigger DSI, identifying DSI as a previously unrecognized pathway linking neuroinflammation to neuronal degeneration (Joshi et al., 2022). However, the relative contributions of synaptic and extra synaptic NMDAR signalling to DSI remain poorly understood. Given the opposing roles of GluN2A- and GluN2B-containing NMDARs in neuronal survival and death, we hypothesized that these receptors differentially regulate DSI under inflammatory conditions.

NMDARs are heterotetrametric glutamate-gated ion channels comprised of GluN1 and GluN2 subunits, whose composition critically determines receptor localization and downstream signalling. GluN2A-containing receptors are predominantly concentrated at synaptic sites and drive signalling pathways associated with neuronal survival, and synaptic plasticity. On the contrary, GluN2B-containing receptors are enriched at extra synaptic regions and preferentially link with pathways involved in mitochondrial dysfunction, oxidative stress, and neuronal death (Hardingham and Bading, 2010; Paoletti et al., 2013; Parsons and Raymond, 2014). Therefore, the balance between synaptic GluN2A and extra synaptic GluN2B signalling is increasingly recognized as a key determinant of neuronal fate in neurodegenerative disorders (Zhou et al., 2013; Ge and Wang, 2023).

In this study, we used primary hippocampal cultures and pharmacological tools to evaluate how synaptic and extrasynaptic NMDAR signaling regulates DSI. We also investigated if adjusting the GluN2A/GluN2B ratio leads to predictable changes in DSI. Our results reveal that DSI acts as a convergence point for excitotoxic and inflammatory pathways. Furthermore, we demonstrate that synaptic and extrasynaptic NMDARs have opposing effects on neuronal vulnerability, shedding new light on the mechanisms driving neurodegeneration in inflammatory MS.

## 2. Materials and Methods

### 2.1. Experimental Animals

Homozygous C57BL/6J mice of either sex were used in this study. All experiments involving animals were performed in accordance with University of Wisconsin–Madison Research Animal Resources and Compliance and approved by the Institutional Animal Care and Use Committee (IACUC) of the University of Wisconsin–Madison (Protocol No. M005922-R02-A01). All mice were housed in groups of maximum 5 in individually ventilated cages (IVC) with sawdust bedding and a nesting material. Animals were maintained on a 12:12h light: dark cycle, with food and water available ad libitum.

### 2.2. Primary cultures of hippocampal neurons

Mouse primary hippocampal neurons were isolated from newborn postnatal day 0 (P0) C57BL6 WT mice. Briefly, P0 mice were decapitated, and hippocampi were isolated in ice-cold high glucose DMEM (Catalog no. 11965092; Gibco) plus 10% fetal bovine serum (FBS) (Catalog no. A5670701; Gibco). The hippocampi were rinsed twice with ice-cold HGDMEM and incubated with Papain (2 mg/ml prepared in HGDMEM) for 30 min at 5% CO_2_ at 37°C. Following papain digestion (2 mg/mL; Catalog No. P4762, Sigma-Aldrich), DNase I (Catalog No. 10104159001, Merck) was added at 2.5 mg/mL for 30 s followed by the addition of 10% FBS to inactivate papain. The digested tissue was gently triturated by pipetting and filtered through a 100 µm sterile cell strainer and spun at 800 rpm for 10 min at room temperature (RT). Cell pellets were resuspended in a 3:1 ratio of neurobasal-B27 and HGDMEM (with 10% fetal bovine serum) and plated on poly-D-lysine-coated (0.2 mg/ml) coverslips at a density of 1.1 × 105 cells/ml. The plated neurons were switched to neurobasal B27 medium (Catalog no. 21103049; Gibco) after 5 h and were treated with 1 μM Ara-C (Catalog no. C1768; Sigma-Aldrich (Merck) after 48 h of plating for 48 h to inhibit the growth of non-neuronal cells in the culture. Cells were then maintained in neurobasal B27 medium with one-third of the media changed every 48 h. Neurons were treated with NMDA (Catalog no. HY-17551; MedChemExpress) and IL-1β (10 ng/mL) for 24 hours and DSI experiments were performed on neurons cultured for 9–12 d.

### 2.3. Pharmacological modulation of synaptic and extrasynaptic NMDA receptor signaling

To investigate the contribution of synaptic and extrasynaptic NMDA receptor (NMDAR) signaling to dendritic syntaphilin intrusion (DSI), cultured hippocampal neurons were pretreated for 30 min with selective pharmacological inhibitors prior to NMDA stimulation (10 μM, 24 h).

To inhibit synaptic NMDAR signaling, neurons were treated with the selective GluN2A antagonist NVP-AAM077 (100 nM) before NMDA exposure. To inhibit extrasynaptic NMDAR signaling, neurons were pretreated with the selective GluN2B antagonist Ro25-6981 (10 μM) before NMDA treatment.

To determine whether the GluN2B–TRPM4 signaling complex regulates DSI, neurons were pretreated with the TRPM4 uncoupling peptides C19 (10 μM) or C8 (10 μM), which disrupt the interaction between GluN2B-containing NMDARs and TRPM4, prior to NMDA stimulation. In separate experiments, TRPM4 channel activity was directly inhibited using TRPM4-In-1 (30 μM) for 30 min before NMDA treatment.

To examine the role of downstream CaMKII signaling, neurons were pretreated with the selective CaMKII inhibitor KN-93 (10 μM) for 30 min before NMDA exposure. Following drug treatments, neurons were fixed and processed for SNPH and MAP2 immunofluorescence, and DSI was quantified as described below.

### 2.4. Lentiviral transduction of GluN2A and GluN2B

To examine the contribution of NMDA receptor subunit composition to dendritic syntaphilin intrusion (DSI), primary hippocampal neurons were transduced with lentiviral vectors expressing either FLAG-tagged GluN2A or FLAG-tagged GluN2B. Neurons were transduced on day 1 in vitro (DIV1) with 200 μL of either Lenti-CMV-FLAG-GluN2A-WT or Lenti-CMV-FLAG-GluN2B-WT virus per culture well (2×10^8 TU/mL), as previously described (Holehonnur et al., 2015). Control cultures received no viral treatment.

Neurons were maintained under standard culture conditions until DIV6, after which they were fixed with 4% paraformaldehyde and processed for immunofluorescence using antibodies against syntaphilin (SNPH), MAP2, and FLAG. DSI was quantified by determining the localization of SNPH-positive puncta within MAP2-positive dendrites using confocal microscopy and ImageJ analysis, as described below.

To verify comparable viral transduction efficiency, FLAG immunofluorescence intensity was quantified in neurons transduced with either GluN2A-FLAG or GluN2B-FLAG lentiviral vectors. FLAG fluorescence was measured in dendrites using ImageJ software, and expression levels were compared between the two groups to confirm equivalent transgene expression.

### 2.5. Immunofluorescence and image acquisition for cultured neurons

For antibody labelling in primary neurons, we fixed coverslips containing neuronal cells with 4% paraformaldehyde (PFA) and 4% sucrose in PBS for 20 min at RT followed by permeabilization with 0.3% Triton X-100 for 15 min and blocking in 10% serum for 1 hr at RT. The neurons were then incubated with primary antibody MAP-2 (Catalog no. AB5622-I; Merck Millipore) (1:500; Millipore), and SNPH (1:500; catalogue #ab192605, Abcam) antibodies overnight at 4°C. The following day these coverslips were washed three times (5 min each) with PBS and incubated with fluorescently conjugated Alexa Fluor secondary antibodies (Alexa Fluor 488 goat anti-rabbit and Alexa Fluor 594 goat anti-mouse; Invitrogen) and mounted on glass slides using Ultra Cruz Aqueous Mounting Medium (catalog no. sc-24941; Santa Cruz Biotechnology). Fluorescent images were acquired using a Nikon A1 confocal microscope with 60× (1.4 numerical aperture) plan apochromatic oil-immersion objectives at 1024 × 1024 resolution. Optical settings were kept identical for all the experimental groups to ensure comparability between experimental groups. Image analysis was accomplished using Nikon Elements.

### 2.6. DSI imaging analysis

Dendritic SNPH localization was quantified by orthogonal confocal analysis. Cultured neurons were stained with MAP2 to identify dendrites and SNPH to identify DSI. To confirm dendritic localization of SNPH immunoreactivity, orthogonal images at three different angles were obtained using NIS-Elements software (Nikon) with a 60× oil lens under a Nikon A1RS Confocal Microscope at the University of Wisconsin–Madison Optical Imaging Core to confirm that SNPH immunoreactivity originates from inside the MAP2-positive dendrites. To quantify DSI, three-dimensional image stacks were analyzed using NIS-Elements software. ImageJ software was used to draw contours of all SNPH immunoreactivity (yellow puncta) within the total MAP2 area (red areas) within the fixed frame. This fixed frame is standardized and applied to all samples. Both areas of yellow and red are measured and recorded by ImageJ and saved in Microsoft Excel. Finally, DSI is computed as the total SNPH area divided by the total MAP2 area for a particular dendritic segment under analysis with Excel. Analysis was performed by one of us who was blinded to the experimental conditions.

### 2.7. Statistics

OriginPro statistical software version 9.4 from Origin Lab and ImageJ with version 1.5v were used to perform the statistical analyses for all the experiments. All data are presented as the mean ± SEM (n = 3 independent experiments). Data were obtained from three independent neuronal cultures (n = 3), with at least 30 dendrites analyzed per experimental group. We have analysed a minimum of N=30 dendrites for three different experimental conditions. Statistical differences in measured variables were assessed between the experimental and control groups by two-tailed unpaired Student’s t-test. One-way Anova followed by Post Hoc Tukey HSD (beta) test was applied when comparison was made among multiple groups. Differences were considered statistically significant at *p* < 0.05.

## 3. Results

### 3.1. Synaptic NMDA receptor blockade markedly enhances DSI

To determine the role of synaptic NMDA receptor signaling in DSI, cultured hippocampal neurons were treated with the GluN2A-selective antagonist NVP-AAM077 prior to NMDA exposure. NMDA treatment alone increased DSI by approximately 320% relative to untreated controls; however, this increase did not reach statistical significance (0.001463±0.000746 vs. 0.007156±0.001194, p=0.1915432). In contrast, inhibition of GluN2A-containing synaptic NMDARs with NVP-AAM077 markedly potentiated NMDA-induced DSI, resulting in an approximately 245% increase compared with NMDA treatment alone (0.007156±0.001194 vs. 0.024175±0.002597, p= 0.0010053; ** p<0.01) and an approximately 1350% increase relative to untreated controls (0.001463±0.000746 vs. 0.024175±0.002597, p= 0.0010053; ** p<0.01) (Figure 1). These findings indicate that synaptic GluN2A-mediated signaling normally suppresses DSI and suggest that physiological synaptic NMDAR activity exerts a protective effect against dendritic SNPH accumulation.

**Fig. 1.**
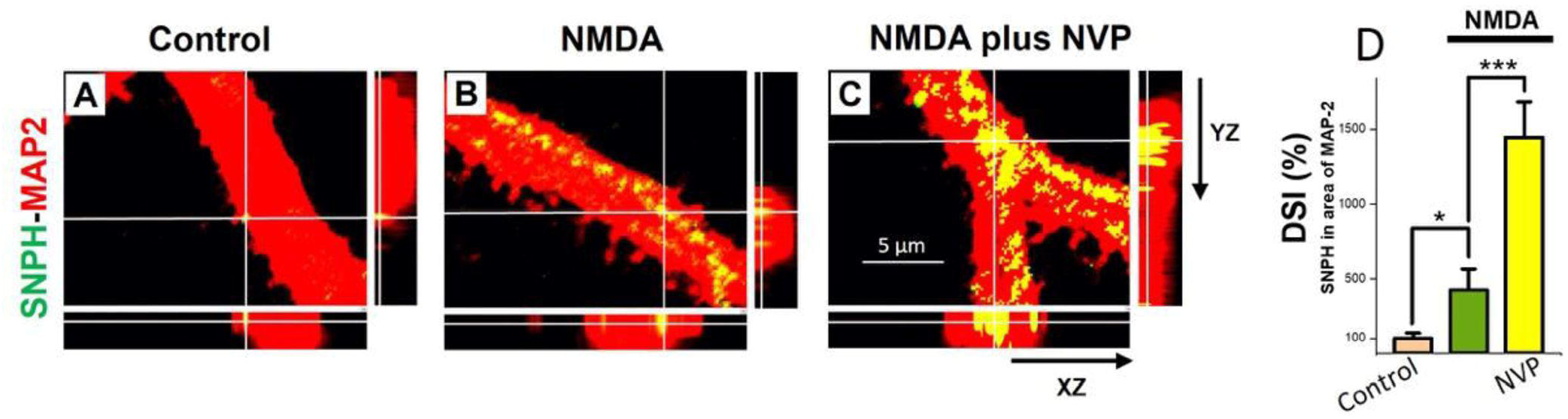
GluN2A is neuroprotective in suppressing DSI: Inhibition of GluN2A greatly increases DSI triggered by NMD. GluN2A is neuroprotective in suppressing DSI. Selective inhibition of GluN2A greatly increases DSI triggered by NMDA. (A – C) Merged images of SNPH-MAP2 staining of cultured hippocampal neurons showing NMDA (10μM, 24 hrs) triggered massive DSI (B) that is tremendously increased by 30 min pre-treatment with selective inhibitor of GluN2A-containing NMDARs, NVP-AAM007 (C, 100 nM); D – Quantification of the stimulatory effects of NVP-AAM007 (100 nM) on DSI induced by NMDA. N=30 dendrites for each group (* p<0.05 between control and NMDA and ***p<0.001 between NMDA and NVP-AAM007).

### 3.2. Inhibition of GluN2B-containing NMDARs suppresses NMDA-induced DSI

To determine whether extrasynaptic GluN2B-containing NMDARs mediate DSI, cultured hippocampal neurons were pretreated with the selective GluN2B antagonist Ro25-6981 before NMDA exposure. NMDA increased DSI by approximately 240% relative to untreated controls (*p* < 0.05). Pretreatment with Ro25-6981 markedly suppressed NMDA-induced DSI, producing an approximately 82% reduction compared with NMDA-treated neurons (*p* < 0.05) and reducing DSI to levels below those observed in untreated controls (Figure 2). These findings demonstrate that activation of GluN2B-containing extrasynaptic NMDARs is required for NMDA-induced dendritic SNPH intrusion.

**Fig. 2.**
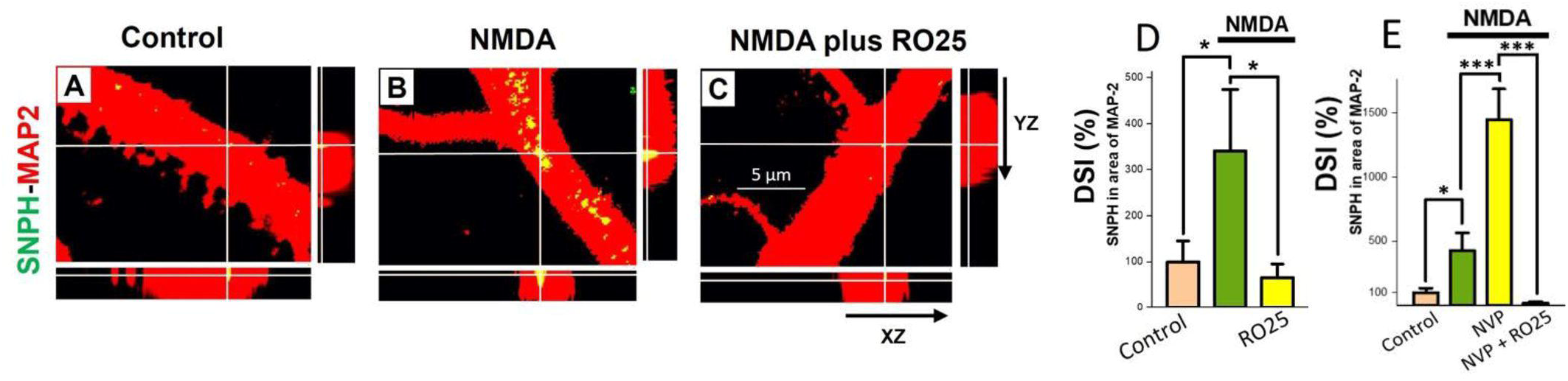
Inhibition of GluN2B greatly prevents DSI triggered by NMDA. GluN2B is neurodegenerative in increasing DSI. Selective inhibition of GluN2B greatly prevents DSI triggered by NMDA. (A – C) Merged images of SNPH-MAP2 staining of cultured hippocampal neurons showing NMDA (10μM, 24 hrs) triggered massive DSI (B) that is blocked by 30 min pre-treatment with selective inhibitor of GluN2B-containing NMDARs, RO25-6981 (C, 10 μM); D – Quantification of the inhibitory effects of tyrosine kinase inhibitor RO25 (10μM) on DSI induced by NMDA. E – DSI increase in NVP is blocked by RO25. N=30 dendrites for each group (* p<0.05 between control and NMDA and *p<0.05 between NMDA and RO25; ***p<0.001 between NVP and NVP+RO25).

To further examine the interplay between GluN2A and GluN2B signaling, DSI was quantified following GluN2A inhibition in the presence or absence of Ro25-6981. Inhibition of GluN2A with NVP-AAM077 increased DSI by approximately 222% relative to NMDA treatment alone and by approximately 1350% compared with untreated controls (*p* < 0.001). Co-treatment with Ro25-6981 dramatically attenuated this effect, reducing DSI by approximately 86%relative to NVP-treated neurons (*p* < 0.001). These findings demonstrate that the pronounced increase in DSI observed following GluN2A inhibition is dependent on GluN2B activation, indicating that GluN2B signaling is the principal driver of dendritic SNPH intrusion.

### 3.3. DSI is selectively driven by GluN2B but not GluN2A

Cultured hippocampal neurons were transduced with lentiviral vectors (2×10^8 TU/mL) expressing either GluN2A or GluN2B to determine whether individual NMDA receptor subunits differentially regulate DSI. Overexpression of GluN2A produced only a modest increase in DSI (approximately 15–20% above control), which was not statistically significant, indicating that increased GluN2A expression alone is insufficient to promote dendritic SNPH intrusion. In contrast, GluN2B overexpression markedly increased DSI by approximately 350–400% relative to control neurons (*p* < 0.05), demonstrating that GluN2B selectively drives DSI formation (Figure 3).

**Fig. 3.**
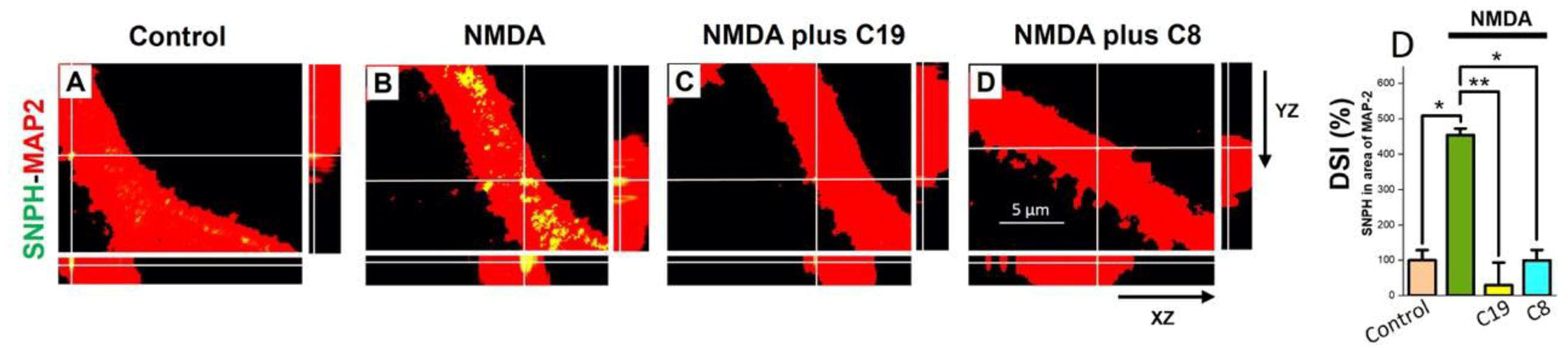
DSI is driven by GluN2B and not GluN2A. Cultured hippocampal neurons were treated with 200 µl of Lenti-CMV Flag-GluN2A Wt and Lenti-CMV Flag-GluN2B Wt virus (2×10^8 TU/mL) on first day of culture and permeabilized and fixed with 4% PFA on sixth day for SNPH, MAP2 and FLAG immunohistochemistry. Ambient DSI unaffected by GluN2A virus (A, C) but dramatically increased by GluN2B virus (B, D). Dendrites N (30 non-virus, 46 GluN2A, 48 for GluN2B). Unpaired t test * p<0.05 between control and GluN2B. (E) No statistical difference (n.s. p=0.09) found between FLAG intensities of GluN2A and GluN2B expression. FLAG intensity measurement in dendrites treated with GluN2A-FLAG or GluN2B-FLAG viruses. FLAG measured in fixed, permeabilized neurons by ImageJ software in dendrites of GluN2A (N=30) and GluN2B (N=28); Unpaired t test applied.

To exclude the possibility that these differences resulted from unequal viral transduction, FLAG immunofluorescence was quantified in neurons expressing GluN2A-FLAG or GluN2B-FLAG. FLAG fluorescence intensity did not differ significantly between the two groups (*p* = 0.09), indicating comparable expression of both constructs. Therefore, the robust increase in DSI observed following GluN2B overexpression reflects a subunit-specific effect rather than differences in transgene expression efficiency.

This experiment demonstrates that GluN2B, but not GluN2A, is sufficient to induce dendritic SNPH intrusion, supporting the conclusion that extrasynaptic GluN2B-containing NMDARs are the principal mediators of DSI.

### 3.4. Disruption of the GluN2B–TRPM4 interaction markedly inhibits NMDA-induced DSI

To determine whether coupling of TRPM4 to GluN2B-containing NMDARs contributes to DSI, cultured hippocampal neurons were pretreated with the TRPM4 uncoupling peptides C19 or C8 before NMDA stimulation. NMDA treatment increased DSI by approximately 350% compared with untreated control neurons (*p* < 0.05). Disruption of the GluN2B–TRPM4 interaction with C19 almost completely abolished NMDA-induced DSI, producing an approximately 93% reduction relative to NMDA-treated neurons (*p* < 0.01) and reducing DSI to levels approximately 70% below control cultures. Similarly, pretreatment with C8 significantly attenuated NMDA-induced DSI by approximately 78% compared with the NMDA group (*p* < 0.05), restoring DSI to values close to those observed in untreated neurons (Figure 4).

**Fig. 4.**
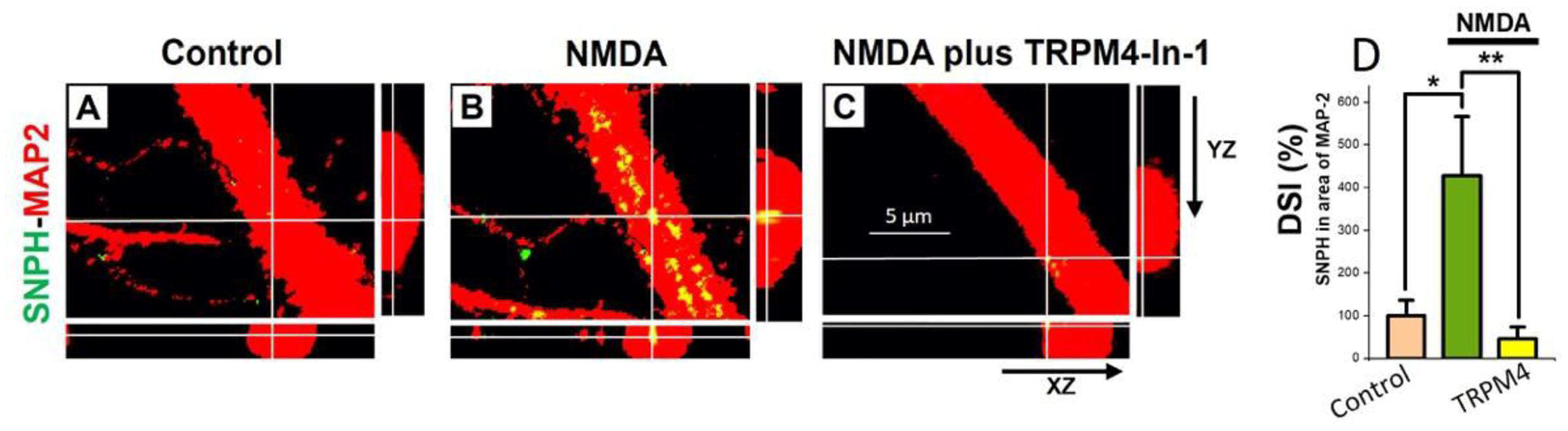
Uncoupling of TRPM4 from GluN2B inhibits DSI triggered by NMDA. The GluN2B partner TRPM4 increases DSI: C19 and C8, uncouplers of TRPM4 from GluN2B, prevent DSI triggered by NMDA (A-D) SNPH-MAP2 merged images showing NMDA (10 μM) triggered DSI (B) blocked by TRPM4 uncouplers C19 (10 μM; C) and C8 (10 μM; D) using orthogonal analysis; E: Quantification of the inhibitory effects of C19 (10μM) and C8 (10μM) on DSI induced by NMDA (10μM). Cells pre-treated with 30 min of C19 and C8 before NMDA was applied for 24 hrs. N=30 dendrites for each group except for NMDA (N=20) and C8 (N=20), One-way Anova with Post Hoc Tukey HSD (beta) test (* p<0.05 between control and NMDA; ** p<0.01 between NMDA and C19; * p<0.05 between NMDA and C8.

These findings demonstrate that the GluN2B–TRPM4 signaling complex is required for NMDA-induced dendritic SNPH intrusion, and that disrupting this interaction effectively prevents DSI.

### 3.5. Direct inhibition of TRPM4 markedly suppresses NMDA-induced DSI

To determine whether TRPM4 activity is required for NMDA-induced DSI, cultured hippocampal neurons were pretreated with the selective TRPM4 inhibitor TRPM4-In-1 before NMDA stimulation. NMDA treatment increased DSI by approximately 325% compared with untreated control neurons (*p* < 0.05), confirming that NMDA receptor activation robustly promotes dendritic SNPH intrusion. In contrast, pharmacological inhibition of TRPM4 with TRPM4-In-1 almost completely abolished NMDA-induced DSI, producing an approximately 89% reduction relative to NMDA-treated neurons (*p* < 0.01). DSI levels in TRPM4-In-1-treated neurons were approximately 55% lower than those observed in untreated controls, indicating that TRPM4 inhibition effectively prevented dendritic SNPH accumulation (Figure 5).

**Fig. 5.**
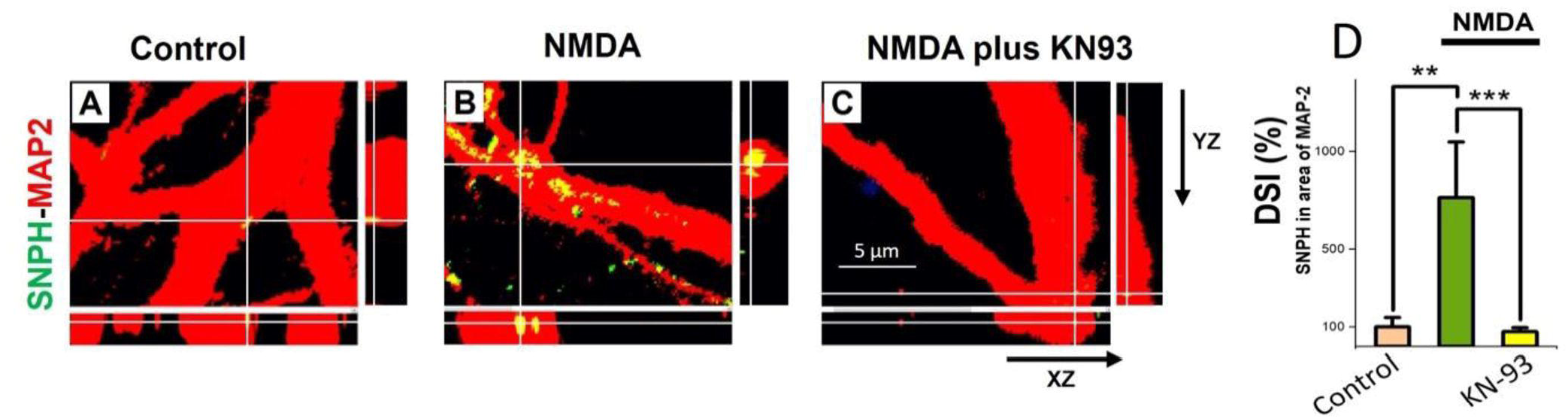
Direct inhibition of TRPM4 inhibits DSI triggered by NMDA. The GluN2B partner TRPM4 increases DSI: Direct inhibition of TRPM4 prevents DSI triggered by NMDA (A-C) SNPH-MAP2 merged images showing NMDA (10 μM) triggered DSI (B) blocked by TRPM4 inhibitor TRPM4-In-1 (30μM; C); D: Quantification of the inhibitory effects of TRPM4-In-1 (30μM) on DSI induced by NMDA (10μM). Cells pre-treated with 30 min of TRPM4-In-1 before NMDA was applied for 24 hrs. N=30 dendrites for each group, One-way Anova with Post Hoc Tukey HSD (beta) test (* p<0.05 between control and NMDA; ** p<0.01 between NMDA and TRPM4-In-1.

These findings demonstrate that TRPM4 activity is essential for NMDA-mediated DSI and further support the conclusion that the GluN2B–TRPM4 signaling axis is a critical upstream regulator of dendritic SNPH mislocalization.

### 3.6. CaMKII inhibition abolishes NMDA-induced DSI

To determine whether CaMKII functions downstream of GluN2B signaling to regulate DSI, cultured hippocampal neurons were pretreated with the selective CaMKII inhibitor KN-93 before NMDA stimulation. NMDA treatment markedly increased DSI by approximately 650% relative to untreated control neurons (p < 0.01), demonstrating robust induction of dendritic SNPH intrusion following NMDA receptor activation. Pharmacological inhibition of CaMKII with KN-93 almost completely prevented NMDA-induced DSI, producing an approximately 96% reduction compared with NMDA-treated neurons (p < 0.001). DSI levels in KN-93-treated neurons were reduced to approximately 70% below those observed in untreated controls, indicating near-complete suppression of dendritic SNPH accumulation (Figure 6).

**Fig. 6.**
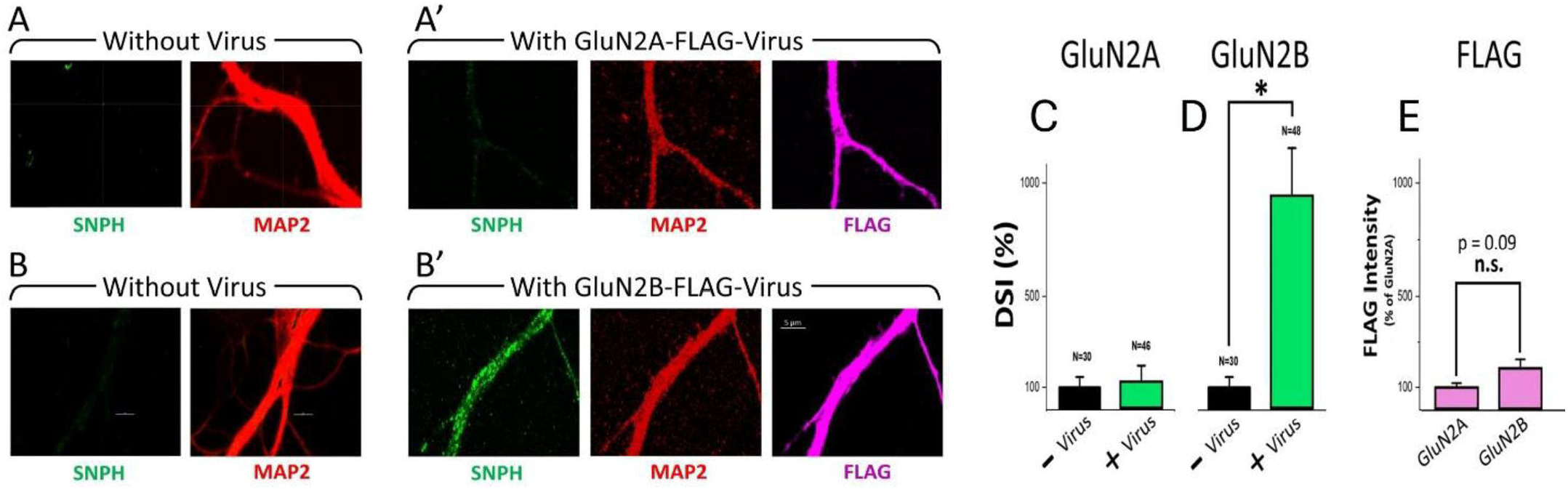
GluN2B interacts with CAMKII: Inhibition of CAMKII prevents DSI triggered by NMDA. (A-C) SNPH-MAP2 merged images showing NMDA (10 μM) triggered DSI (B) blocked by CAMKII inhibitor KN93 (10μM; C); D: Quantification of the inhibitory effects of KN93 (10μM) on DSI induced by NMDA (10μM). Cells pre-treated with 30 min of KN93 before NMDA was applied for 24 hrs. N=30 dendrites for each group, One-way Anova with Post Hoc Tukey HSD (beta) test (** p<0.01 between control and NMDA; *** p<0.001 between NMDA and KN93)

These findings demonstrate that CaMKII activation is essential for NMDA-induced DSI and place CaMKII downstream of GluN2B–TRPM4 signaling in the molecular pathway regulating dendritic SNPH intrusion.

### 3.7. DSI is a downstream convergence point for synaptic and extrasynaptic NMDA receptor signaling

The collective findings support a model in which synaptic and extrasynaptic NMDA receptors exert opposing influences on neuronal survival through regulation of DSI. Synaptic GluN2A-containing receptors suppress DSI and maintain dendritic mitochondrial homeostasis, whereas extrasynaptic GluN2B-containing receptors activate TRPM4- and CaMKII-dependent signaling pathways that promote SNPH mislocalization and dendritic pathology. Viral enhancement of GluN2B signaling further sensitizes neurons to DSI, while blockade of extrasynaptic pathways protects against its development.

These results identify DSI as a previously unrecognized downstream target of the GluN2A/GluN2B signaling balance and establish DSI as a mechanistic convergence point linking inflammatory cytokines, excitotoxic signaling, mitochondrial dysfunction, and neurodegeneration in multiple sclerosis (Figure 7).

**Fig. 7.**
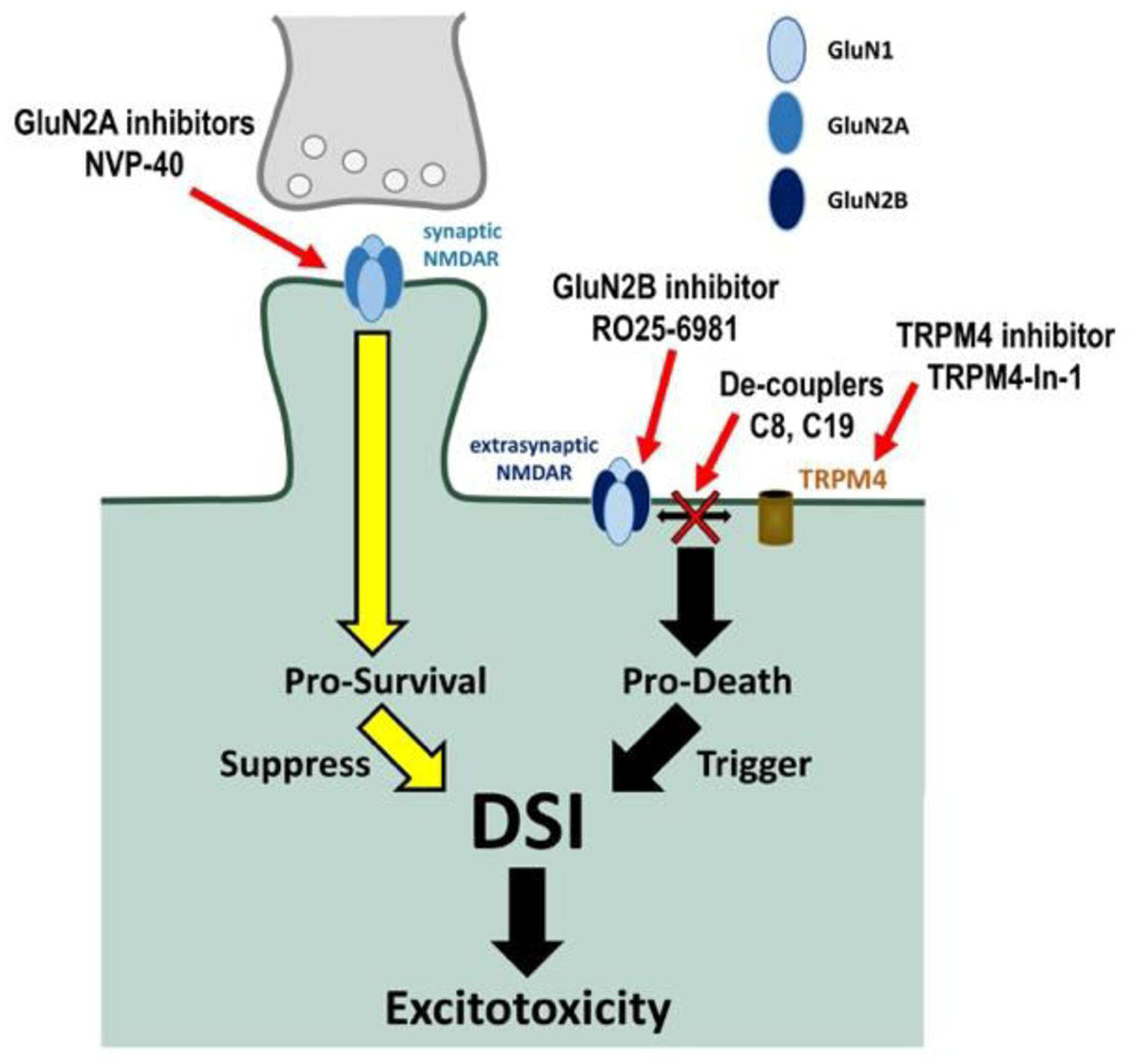
Summary showing opposite regulation of DSI by synaptic and extrasynaptic NMDA receptors.

## Discussion

The present study identifies Dendritic Syntaphilin Intrusion (DSI) as a novel downstream mechanism through which inflammatory and excitotoxic signaling converge to promote neurodegeneration in Multiple Sclerosis (MS). Although glutamate-mediated excitotoxicity has long been implicated in MS pathology, the molecular pathways linking neuroinflammation to dendritic dysfunction remain incompletely understood (Centonze et al., 2010; Stojanovic et al., 2014). Our findings demonstrate that the balance between synaptic and extrasynaptic NMDA receptor signaling is a critical determinant of DSI and that opposing actions of GluN2A- and GluN2B-containing receptors regulate neuronal vulnerability. Specifically, inhibition of synaptic NMDA receptors enhanced DSI, whereas blockade of extrasynaptic NMDA receptors suppressed DSI. Furthermore, viral overexpression of GluN2B markedly increased ambient DSI, while GluN2A overexpression failed to induce a similar effect. Together, these findings identify DSI as a downstream target of the GluN2A/GluN2B signaling balance and establish a direct mechanistic connection between neuroinflammation, excitotoxicity, mitochondrial dysfunction, and grey matter degeneration.

### DSI as a novel mitochondrial mechanism of neurodegeneration

Mitochondrial dysfunction is increasingly recognized as a major contributor to neurodegeneration in progressive MS (Dutta and Trapp, 2011; Mahad et al., 2015). Proper mitochondrial distribution within neuronal processes is essential for maintaining local energy supply and calcium buffering. Syntaphilin (SNPH) normally functions as an axonal mitochondrial anchor that immobilizes mitochondria at sites of high metabolic demand (Kang et al., 2008). In contrast, dendritic mitochondria remain highly dynamic to support synaptic plasticity and neuronal adaptability (Chen and Sheng, 2013; Devine and Kittler, 2018).

Our previous work demonstrated that aberrant translocation of SNPH into dendrites immobilizes dendritic mitochondria and initiates a novel form of excitotoxic injury termed DSI (Joshi et al., 2019). The present study extends these observations by demonstrating that DSI is regulated by inflammatory and excitotoxic pathways directly relevant to MS. These findings suggest that DSI represents a previously unrecognized mitochondrial mechanism contributing to grey matter pathology and neuronal dysfunction.

### Opposing roles of synaptic and extrasynaptic NMDA receptor signaling

A major finding of this study is the opposing regulation of DSI by synaptic and extrasynaptic NMDA receptor populations. Pharmacological inhibition of GluN2A-containing synaptic NMDA receptors significantly enhanced DSI, indicating that physiological synaptic signaling suppresses dendritic mitochondrial pathology. In contrast, inhibition of GluN2B-containing extrasynaptic NMDA receptors markedly reduced DSI, demonstrating that extrasynaptic signaling is a principal driver of SNPH mislocalization.

These observations are consistent with the well-established dichotomy of NMDA receptor signaling. Synaptic NMDA receptors activate pro-survival pathways involving CREB, BDNF, and activity-dependent gene expression, whereas extrasynaptic receptors initiate cell-death pathways associated with mitochondrial dysfunction, oxidative stress, and neurodegeneration (Hardingham and Bading, 2010; Parsons and Raymond, 2014; Zhou et al., 2013; Ge and Wang, 2023). The dramatic increase in ambient DSI following GluN2B overexpression further supports the concept that receptor composition strongly influences neuronal susceptibility to degeneration. Collectively, these findings identify the GluN2A/GluN2B ratio as a critical regulator of DSI and neuronal survival.

The opposing effects of GluN2A and GluN2B signaling observed in the present study are consistent with the broader framework of NMDA receptor compartmentalization. Synaptic NMDAR activation promotes CREB-dependent transcription, mitochondrial biogenesis, and neuronal resilience, whereas extrasynaptic NMDAR activation suppresses survival pathways and induces pro-degenerative signaling cascades (Hardingham and Bading, 2010; Paoletti et al., 2013). Our findings extend this concept by identifying DSI as a previously unrecognized downstream effector through which NMDAR subtype-specific signaling influences neuronal vulnerability. Thus, DSI may represent a mechanistic bridge linking receptor localization to mitochondrial dysfunction and neurodegeneration.

### GluN2B–TRPM4–CaMKII signaling drives DSI

The present study further identifies downstream signaling mechanisms linking GluN2B activation to SNPH mislocalization. The neuroprotective effects observed following GluN2B inhibition are consistent with previous studies demonstrating beneficial effects of GluN2B-selective antagonists in experimental neurodegeneration and neuroinflammatory disease models (Taniguchi et al., 1997; Farjam et al., 2014; Quillinan et al., 2015; Mares et al., 2021). Both pharmacological inhibition of TRPM4 and disruption of the GluN2B–TRPM4 interaction significantly reduced NMDA-induced DSI. These findings are consistent with previous reports demonstrating that the GluN2B–TRPM4 complex functions as a critical mediator of excitotoxic signaling (Yan et al., 2020).

In addition, inhibition of CaMKII suppressed DSI, indicating that CaMKII operates downstream of GluN2B signaling. Given the established association between GluN2B and CaMKII in excitotoxic pathways (Strack and Colbran, 1998; Tullis et al., 2021), our findings support the existence of a GluN2B–TRPM4–CaMKII signaling axis upstream of SNPH mislocalization. This pathway provides a mechanistic framework linking excitotoxic receptor activation to mitochondrial dysfunction and dendritic degeneration.

### Relevance to inflammatory multiple sclerosis

An important implication of the present findings is the extension of DSI from non-inflammatory models to inflammatory MS. Pro-inflammatory cytokines such as IL-1β and TNF-α are elevated in active MS lesions and are known to potentiate glutamatergic excitotoxicity by enhancing extrasynaptic NMDA receptor signaling while diminishing neuroprotective synaptic activity (Viviani et al., 2003; Mandolesi et al., 2013; Rossi et al., 2014). Our previous work demonstrated reciprocal interactions between IL-1β and NMDA receptor activation in triggering DSI (Joshi et al., 2022).

The current study suggests that inflammatory cytokines facilitate DSI by shifting NMDA receptor signaling toward GluN2B-dependent pathways. Consequently, DSI may represent a critical downstream mechanism through which neuroinflammation is translated into progressive neuronal dysfunction and grey matter degeneration. This concept is particularly relevant to progressive MS, where grey matter pathology strongly correlates with disability progression (Calabrese et al., 2015; Klaver et al., 2013; Zhang et al., 2021).

### Implications for other neurodegenerative diseases

The significance of DSI may extend beyond MS. Dysregulation of synaptic and extrasynaptic NMDA receptor signaling has been implicated in Alzheimer’s disease, Huntington’s disease, amyotrophic lateral sclerosis, traumatic brain injury, and ischemic neurodegeneration (Hardingham and Bading, 2010; Lau and Tymianski, 2010; Fairless et al., 2021). Because DSI occurs downstream of these pathways, aberrant SNPH localization may represent a common mitochondrial mechanism contributing to neuronal degeneration across diverse neurological disorders. Future studies should determine whether DSI is present in these disease contexts and whether targeting SNPH can provide broad neuroprotective benefits.

### Therapeutic implications

The opposing effects of GluN2A and GluN2B signaling suggest that restoration of NMDA receptor subtype balance may represent a rational therapeutic strategy. Unlike global NMDA receptor blockade, which disrupts physiological synaptic transmission, selective enhancement of GluN2A-mediated signaling combined with suppression of GluN2B-dependent pathways could preserve beneficial neuronal functions while limiting excitotoxic injury (Hardingham and Bading, 2010; Rossi et al., 2013).

Importantly, SNPH itself may represent a therapeutic target because it functions downstream of multiple inflammatory and excitotoxic pathways. Targeting DSI could therefore provide a strategy for preventing mitochondrial dysfunction and neuronal degeneration regardless of the initiating insult (Joshi et al., 2019; Joshi et al., 2022).

## Limitations

There are several limitations of this study. First, the majority of experiments were performed in primary hippocampal neuronal cultures, which do not fully recapitulate the complex inflammatory and cellular environment of MS lesions. Second, although the identified signaling pathways are highly relevant to inflammatory MS, direct evidence of DSI in human MS tissue remains unavailable. Third, the current study focused primarily on NMDA receptor signaling and did not investigate the potential contributions of additional excitotoxic pathways, including AMPA receptor signaling, oxidative stress, and microglial-mediated mechanisms. Finally, while viral manipulation of GluN2A and GluN2B strongly supports a causal role for receptor subtype balance, in vivo validation in inflammatory MS models such as EAE will be essential to establish the physiological significance of DSI during disease progression.

## Conclusions

In conclusion, this study demonstrates that Dendritic Syntaphilin Intrusion (DSI) is oppositely regulated by synaptic and extrasynaptic NMDA receptors and DSI may represent an important mechanism contributing to neuroinflammation, excitotoxicity, mitochondrial dysfunction, and neurodegeneration. Synaptic GluN2A-containing NMDA receptors suppress DSI and promote neuronal survival, whereas extrasynaptic GluN2B-containing receptors drive DSI through TRPM4- and CaMKII-dependent signaling pathways. These findings extend the relevance of DSI to inflammatory MS and establish the GluN2A/GluN2B balance as a critical determinant of neuronal vulnerability. Targeting DSI, either through restoration of synaptic-extrasynaptic NMDA receptor balance or direct modulation of SNPH-mediated mitochondrial anchoring, may represent a promising therapeutic strategy for MS and other neurodegenerative disorders.

## Acknowledgement

This work was supported by NIH R01 MSN233781 granted to SY Chiu. We thank Lance Rodenkirch (University of Wisconsin–Madison Optical Imaging Core) for fluorescence imaging support. We thank Professor Jonathan E. Ploski, Department of Neuroscience and Experimental Therapeutics, Penn State College of Medicine, Hershey, PA, USA, for generously providing the virus.

## Author Contributions

S.Y.C. designed research; D.M., and C.L.Z. performed experiments, analysed data, contributed to data acquisition and preparing figures; D.M., C.L.Z. and S.Y.C. wrote the paper.

## Potential Conflicts of Interest

The authors declare no competing financial interests.

## Data Availability

The data that support the findings of this study are available from the corresponding authors, upon reasonable request.

## Notes

### Competing Interest Statement

The authors have declared no competing interest.

